# Microbial communities profiling in intensive care units expose limitations in current sanitary standards

**DOI:** 10.1101/633404

**Authors:** Lucas Ferreira Ribeiro, Erica M. Lopes, Luciano T. Kishi, Liliane Fraga Costa Ribeiro, Mayra Gonçalves Menegueti, Gilberto Gambero Gaspar, Rafael Silva-Rocha, María Eugenia Guazzaroni

## Abstract

Hospital-associated infections (HAIs) are the leading cause of morbidity and mortality in intensive care units (ICUs) and neonatal intensive care units (NICUs). Organisms causing these infections are often present on surfaces around the patient. Given that microbiotas may vary across different ICUs, the HAI-related microbial signatures within these units remain underexplored. In this study, we use deep-sequencing analyses to explore and compare the structure of bacterial communities at inanimate surfaces of the ICU and NICU wards of The Medical School Clinics Hospital (Brazil). The data revealed that NICU presents higher biodiversity than ICU and surfaces closest to the patient showed a peculiar microbiota, distinguishing one unit from the other. Several facultative anaerobes or obligate anaerobes HAI-related genera were classified as biomarkers for the NICU, whereas *Pseudomonas* was the main biomarker for ICU. Correlation analyses revealed a distinct pattern of microbe-microbe interactions for each unit, including bacteria able to form multi-genera biofilms. Furthermore, we evaluated the effect of concurrent cleaning over the ICU bacterial community. The results showed that, although some bacterial populations decreased after cleaning, various HAI-related genera were quite stable to sanitization, suggesting being well-adapted to the ICU environment. Overall, these results enabled identification of discrete ICU and NICU reservoirs of potentially pathogenic bacteria and provided evidence for the presence of a set of biomarkers that distinguish these units. Moreover, the study exposed the inconsistencies of the routine cleaning to minimize HAI-related genera contamination.

**IMPORTANCE:** Due to the high impact of HAIs, there is an urgent need for the development of robust policies on microbial surveillance to help guide procedures, improving infection control. To the best of our knowledge, this is the first comprehensive study, using a high-throughput approach, focused on comparing the microbiota peculiarities of the ICU and NICU in one of the largest public hospitals in Brazil. The work highlighted bacteria associated with nosocomial infections, identifying the most potent reservoirs of contamination, and evaluated the microbiota changes related to the cleaning procedure. Therefore, this study contributes to increase the knowledge about (N)ICUs microbiomes and may help to reduce health-care-associated infections, especially in developing countries.

## INTRODUCTION

Microbiome refers to the microbial community, and their respective genomes, associated with a particular habitat, including natural or built environments [1]. Natural ecosystems have been well explored; however, not much is known about indoor microbiomes – offices, houses, buildings, hospitals, etc. – where the majority of our life is spent and can have a severe impact on human health. Unlike most indoor environments, intensive care units (ICUs) or neonatal intensive care unit (NICUs) in hospitals are routinely monitored by standard cultivation techniques [2,3]. Nonetheless, conventional cultivation techniques can identify only a tiny proportion of the total bacteria [2,4]. Oberauner et al. [2] reported that only 2.5% of the overall bacterial diversity were identified in an ICU microbiome using culture-dependent methods. Culture-independent methods such as next-generation sequencing (NGS) technologies have a tremendous effect on profiling microbiomes. Phylogenetic analyses based on 16S gene diversity have been fundamental to uncover (N)ICU bacterial varieties in depth and at high resolution in space and time, and it can contribute to improving hospital safety.

In (N)ICUs, even adopting strict sanitation protocols, many patients are infected with healthcare-associated infections (HAIs), also known as nosocomial infections, a significant public health problem around the world [5–10]. HAIs include diseases that can be associated with surfaces and devices present in hospitals and can spread through health care staff, contaminated surfaces or air droplets. These infections are more frequent in UTIs where outbreaks often originate [11]. HAIs increase deaths (morbidity and mortality), antimicrobial resistance, prolong the duration of hospital stays, and consequentially healthcare costs [12]. The National Healthcare Safety Network of the Centers for Disease Control and Prevention (CDC) has estimated 687,000 HCAIs in U.S. acute care hospitals causing 72,000 deaths, and costs estimated to $97-147 billion annually [13,14]. The most common pathogen causing HAIs are *Clostridium difficile* and ‘ESKAPE’ bacteria (*Enterococcus spp*., *Staphylococcus aureus*, *Klebsiella spp*., *Acinetobacter spp*., *Pseudomonas aeruginosa*, and Enterobacteriaceae) [14,15]. Many of these bacteria exhibit antimicrobial resistance and can cause infections of the bloodstream, urinary tract, severe pneumonia, and surgical site infection [11,16].

Hospital surfaces remain neglected reservoirs for HAI-related bacteria, and strict cleaning protocols have been used as the primary procedure to reduce the risks. Nonetheless, the efficiency of cleaning protocols, usually, has been investigated by culture-dependent routine techniques. Here, using NGS methodology, we analyzed the differences and similarities between the structure of bacterial communities from the ICU and NICU surfaces of The Medical School Clinics Hospital (Ribeirão Preto, Brazil), one of the biggest hospitals in Latin America, and which has more than 35,000 hospitalizations per year and supports a population of four million people. We hypothesized that the microbiota “signature” would vary significantly between ICU and NICU, offering opportunities for targeted spatial biomarkers to improve the combat against HAIs. Furthermore, we tested the impact of the standard cleaning procedure established on the hospital on ICU microbiota, focusing on bacteria associated with nosocomial infections.

## RESULTS AND DISCUSSION

### Microbial profiling of ICU and NICU samples using V4-5 16 rRNA sequencing

In order to compare the microbial community of the ICU and NICU from a clinical hospital in Brazil, we use NGS targeting V4 hypervariable regions within microbial 16S rRNA genes [17]. The intensive care units contained two wards with four beds each (Fig.1), where critically ill patients were present. Samples were collected from boxes areas (mattresses, bed rails, monitors, infusion pumps, ventilators, and cufflator), with patients lying down; and also, in common areas (computers-keyboard and mouse, doors handle, hospital cards, medical records, drug stations, and nurse’s mobiles). Furthermore, to address the question of how concurrent cleaning impacts the microbial ecosystem of an ICU, samples were sequenced either before or immediately after cleaning.

**Figure 1.**
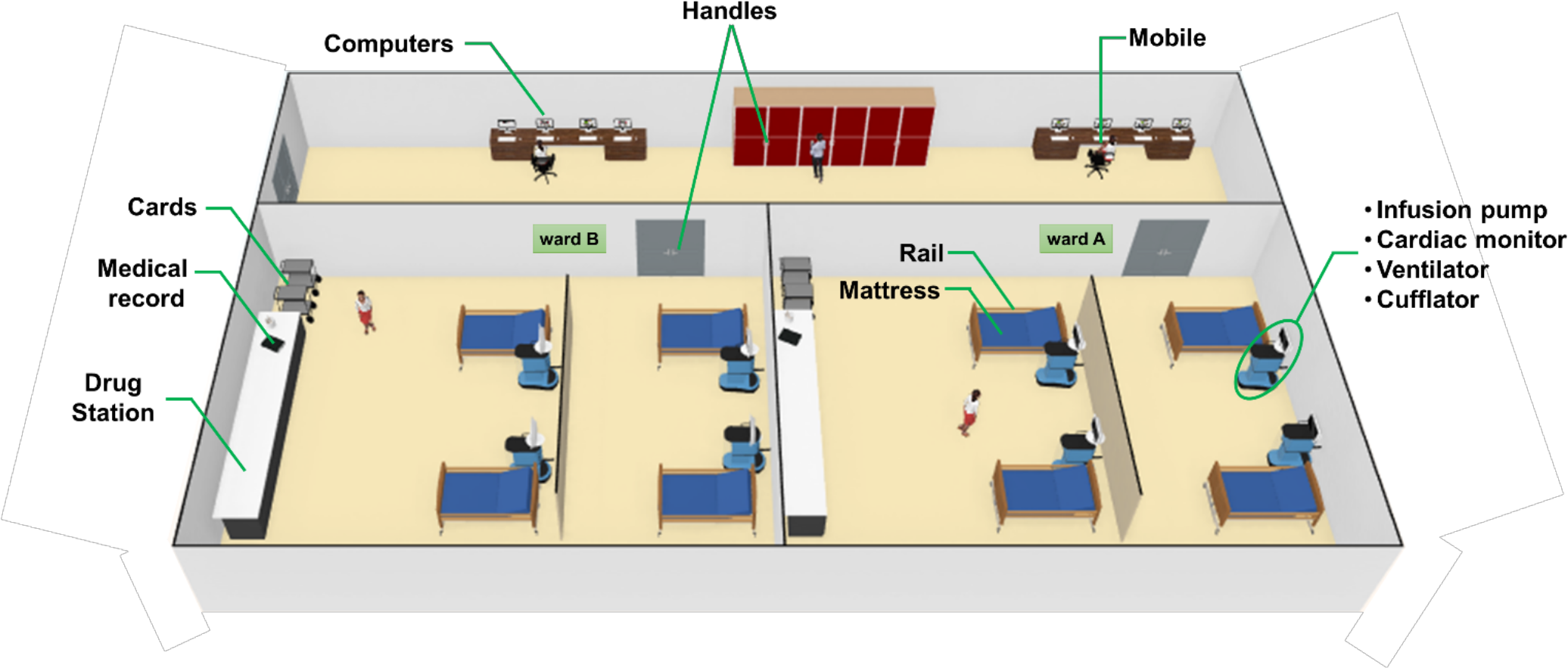
3D-rendered-model showing each sampling site of the (Neonatal) intensive care unit ((N)ICU). ICU and NICU are on different floors at the hospital but have a similar arrangement of wards and devices in general. A detailed explanation of each sample is shown in Table 1.

**Table 1.** Essential characteristics and localization of the sequenced samples.

| Sample ID | Sample source | Care unit | Ward | Cleaning |
| --- | --- | --- | --- | --- |
| Pumps-ICUa | Pump | ICU | A | — |
| Mattresses-ICUa | Mattress | ICU | A | — |
| Rails-ICUa | Rail | ICU | A | — |
| Monitors-ICUa | Monitor | ICU | A | — |
| Ventilators-ICUa | Ventilator | ICU | A | — |
| Pumps-ICUb | Pump | ICU | B | — |
| Mattresses-ICUb | Mattress | ICU | B | — |
| Rails-ICUb | Rail | ICU | B | — |
| Monitors-ICUb | Monitor | ICU | B | — |
| Ventilators-ICUb | Ventilator | ICU | B | — |
| MedicalRecords-ICUab | Medical record | ICU | AB | — |
| Cards-ICUab | Card | ICU | AB | — |
| Mobiles-ICUab | Mobiles | ICU | AB | — |
| Handles-ICUab | Handle | ICU | AB | — |
| DrugStations-ICUab | Drug station | ICU | AB | — |
| Computers-ICUab | Computer | ICU | AB | — |
| Pumps-ICUaA | Pump | ICU | A | + |
| Mattresses-ICUaA | Mattress | ICU | A | + |
| Rails-ICUaA | Rail | ICU | A | + |
| Monitors-ICUaA | Monitor | ICU | A | + |
| Ventilators-ICUaA | Ventilator | ICU | A | + |
| Pumps-ICUbA | Pump | ICU | B | + |
| Mattresses-ICUbA | Mattress | ICU | B | + |
| Rails-ICUbA | Rail | ICU | B | + |
| Monitors-ICUbA | Monitor | ICU | B | + |
| Ventilators-ICUbA | Ventilator | ICU | B | + |
| Cufflators-ICUabA | Cufflator | ICU | AB | + |
| Pumps-NICUa | Pump | NICU | A | — |
| Mattresses-NICUa | Mattress | NICU | A | — |
| Rails-NICUa | Rail | NICU | A | — |
| Monitors-NICUa | Monitor | NICU | A | — |
| Ventilators-NICUa | Ventilator | NICU | A | — |
| Pumps-NICUb | Pump | NICU | B | — |
| Mattresses-NICUb | Mattress | NICU | B | — |
| Rails-NICUb | Rail | NICU | B | — |
| Monitors-NICUb | Monitor | NICU | B | — |
| Ventilators-NICUb | Ventilator | NICU | B | — |
| Mobiles-NICUab | Mobiles | NICU | AB | — |
| Cards-NICUab | Card | NICU | AB | — |
| Handles-NICUab | Handle | NICU | AB | — |
| MedicalRecords-NICUab | Medical record | NICU | AB | — |
| DrugStations-NICUab | Drug station | NICU | AB | — |
| Computers-NICUab | Computer | NICU | AB | — |

A total of ~1.7 million sequences corresponding to 4.94 Gbp of data from 44 samples were generated. The average number of read counts per sample was 34.621, ranging from 33.708 to 34.739. Thus, the data counts were normalized to 33.708 reads. After trimming, the final number of operational taxonomic unit (OTU) consisted of 2054, 1586, OTUs for NICU, and ICU, respectively. Rarefaction curves (Fig. S1) based on the number of OTUs observed were comparably close to asymptotic for all samples. The cut-off was set to 10,000 sequences per sample whereby the rarefaction curves of all samples reached saturation, indicating the availability of enough covering to represent and compare the microbiome community present within the samples. Chimera and singleton OTU removal was included in the data processing pipeline to prevent overestimated richness. Bellow, we presented the analysis regarding the microbial composition for each sample and the comparison between the different areas analyzed.

### Comparative assessment between ICU and NICU microbiota

Microbial profiling of the ICU and NICU allowed the identification of nine different bacterial phyla: *Firmicutes*, *Proteobacteria*, *Actinobacteria*, *Bacteroidetes*, *Fusobacteria*, *Cyanobacteria*, *Deinococcus*, *Gemmatimonadetes*, and *Euryarchaeota*, while this last was only found in NICU. *Firmicutes* and *Proteobacteria* were the most abundant phyla across all samples, composing 46% and 39% of these bacterial communities, respectively. The over-representation of these phyla agree with previous results obtained for microbial communities found in (N)ICUs inanimate surfaces [2,18–20]. The microbial communities at the genus level (Fig. 2A) included sequences of 138 and 160 genera, for ICU and NICU respectively, among which a substantial number of organisms are not culturable. For all samples, the relative abundance of Not_Assigned (NA) genera was notably moderated (up to 18%). Gram-positive bacteria were found in higher abundance in both units. Nonetheless, in terms of the number of genera, Gram-negative bacteria were more diverse. The number of strictly aerobic genera were highly represented (50%) followed by facultative anaerobe (36%) and obligatory anaerobic bacteria (14%) for both units (see details in supplemental material). *Bacillus*, *Staphylococcus*, and *Pseudomonas* were the most abundant genera (47% of the total reads) on ICU surfaces, and *Bacillus*, *Propionibacterium* and *Staphylococcus* predominated in NICU (40%). These genera contain many commensal species for humans, although it also includes members associated with nosocomial infections in (N)ICUs. Members from these genera are considered “survival specialists,” and can persist for months on dry surfaces [21] or associated with spore or biofilm formation [22,23]. A total of 110 OTUs were found only in ICU and 578 only in NICU, while 1476 OTUs were shared between the units (Fig. S2A).

**Figure 2.**
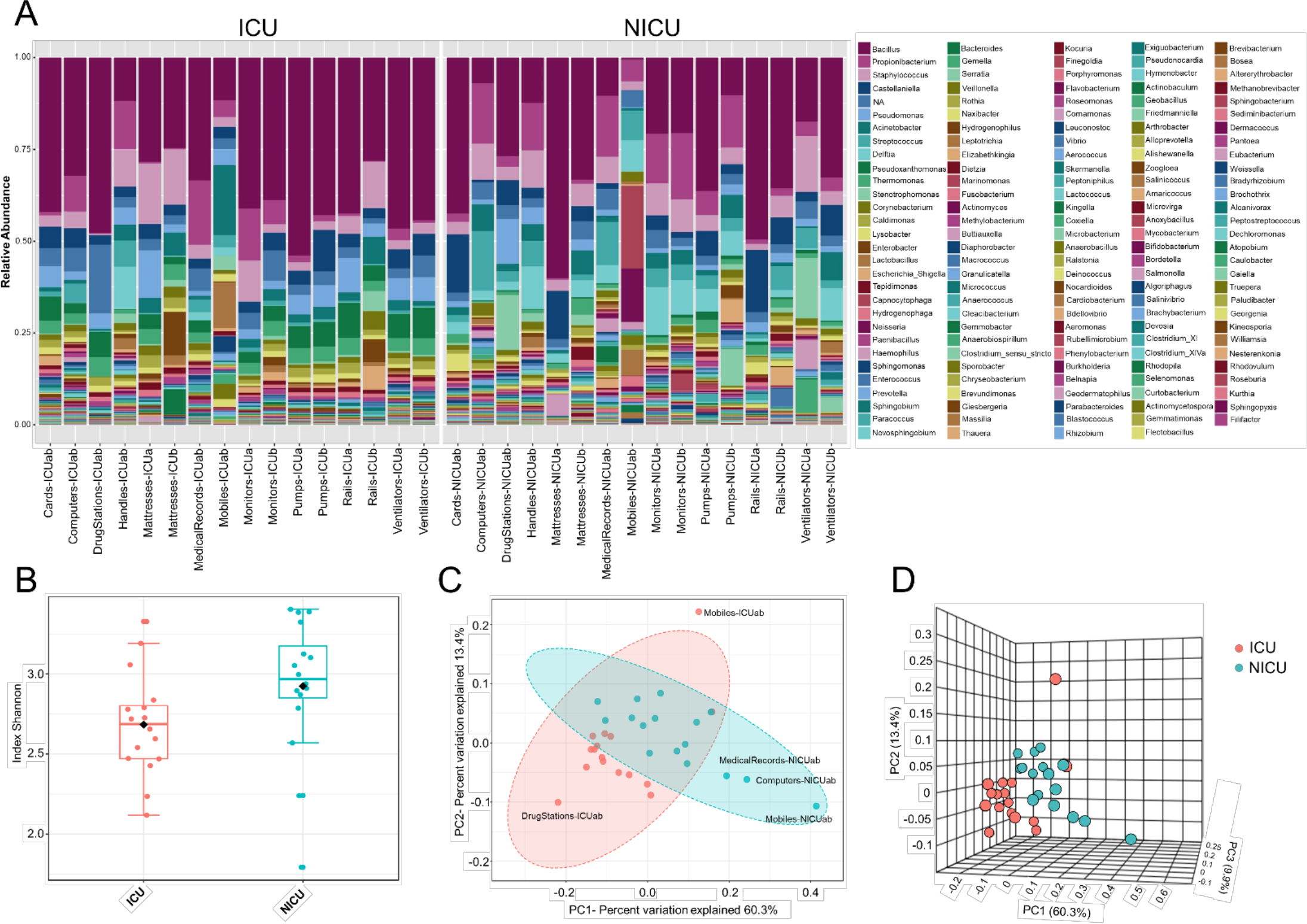
The ICU and NICU bacteria microbiota profile. (**A**) The relative abundance of bacterial genera within the top 379 OTUs among the two units. Colors correspond to the bacterial genera in the legend. Rectangles represent specific genera organized in order of abundance. Sequencing results are presented for each sample clustered using Usearch algorithm with a 97% cutoff. NA (Not_Assigned) represents sequences reads that were not assigning an accurate taxonomic label at the genus level but assigned at the higher taxonomic level. (**B**) Alpha diversity at OTU level at ICU (red, n=16) and NICU (cyan, n=16) calculated using Shannon index (Kruskal-Wallis test, p-value < 0.05). For each box plot herein forward, the line within the box and the black diamond represent the median and mean, respectively. The bottom and top boundaries of each box indicate the first and third quartiles (the 25th and 75th percentiles), respectively. The whiskers represent the lowest and highest values within the 1.5 interquartile range (IQR). Two- (**C**) and three-dimensional (**D**) principal coordinate analysis (PCoA) plot based on Jensen-Shannon distances between bacterial communities associated with ICU and NICU areas (ANOSIM, R= 0.3066; p-value < 0.001). Samples are shown as single dots. Divergence at OTU level was computed on Total sum scaling–normalized (TSS-normalized) datasets.

Analysis of all samples from the care units indicated that NICU samples showed a significantly higher Shannon index – a measure of diversity – as compared to samples belonging to ICU (Kruskal-Wallis test, p-value < 0.05) (Fig. 2B). However, noticeable variation was observed within the sample types (Fig. S2B), and computers and doors handle from both units showed the highest diversity among all samples. A higher Shannon index for NICU agrees with the differences in the number of OTUs found in the care units. The greater diversity in NICU could be explained, in part, due to the higher transit of visitors (e.g., children’s parents or relatives) compared with the more restrictive transit in ICU.

Beta diversity analysis (Fig. 2C-D) of the microbiota for each care unit revealed distinct, but overlapping, profile (ANOSIM, R= 0.3066; p-value < 0.001). A high level of variation among some samples was observed supplemented by less pronounced but distinct variation between ICU/NICU samples closer to the patient (boxes area) (ANOSIM, R= 0.50756; p-value < 0.001) (Fig. S2C). Samples from the common area did not show a significant difference (Fig. S2D). Boxes area samples from ward A and B, belonging to the same care unit, did not show a significant difference (Fig. S2E-F). This analysis suggests that ICU and NICU carry a distinct microbial diversity. Besides, it is also important to remark that more significant differences were observed in the confined area closer to the patients (boxes). These areas are selective environments, where antimicrobial therapies and stringent cleaning protocols are routinely applied.

### Identification of HAI-related genera in neglected (N)ICU surfaces

Evidence suggests that hospital computers (keyboard and mouse) and staff’s mobiles may serve as reservoirs for bacteria associated with HAI within the healthcare environment and facilitate the cross-contamination among hospital wards [24–26]. Taxonomically, ICU mobiles revealed a far greater abundance of *Acinetobacter, Sphingomonas*, and *Brevundimonas* (Fig. 3A). These genera are usually found in moist environments and can show a high risk for HAI in immunocompromised patients. Besides, other genera associated with human microflora were also found in high abundances, such as *Lactobacillus* (mouth and vaginal flora) and *Anaerobiospirillum* (human, cat, and dog feces) [27]. NICU mobiles showed a greater abundance of *Fusobacterium, Neisseria, Rothia, Granulicatella, and Streptococcus* (Fig. 3A) that are part of the oronasopharynx or skin microflora. However, they can also be associated with severe infections in patients with a weakened immune system. Our data are consistent with previous studies that have reported that although mobiles can work as a repository to opportunistic pathogens, portions of their bacteria are also found on the human microbiome (owner’s body) [28].

**Figure 3.**
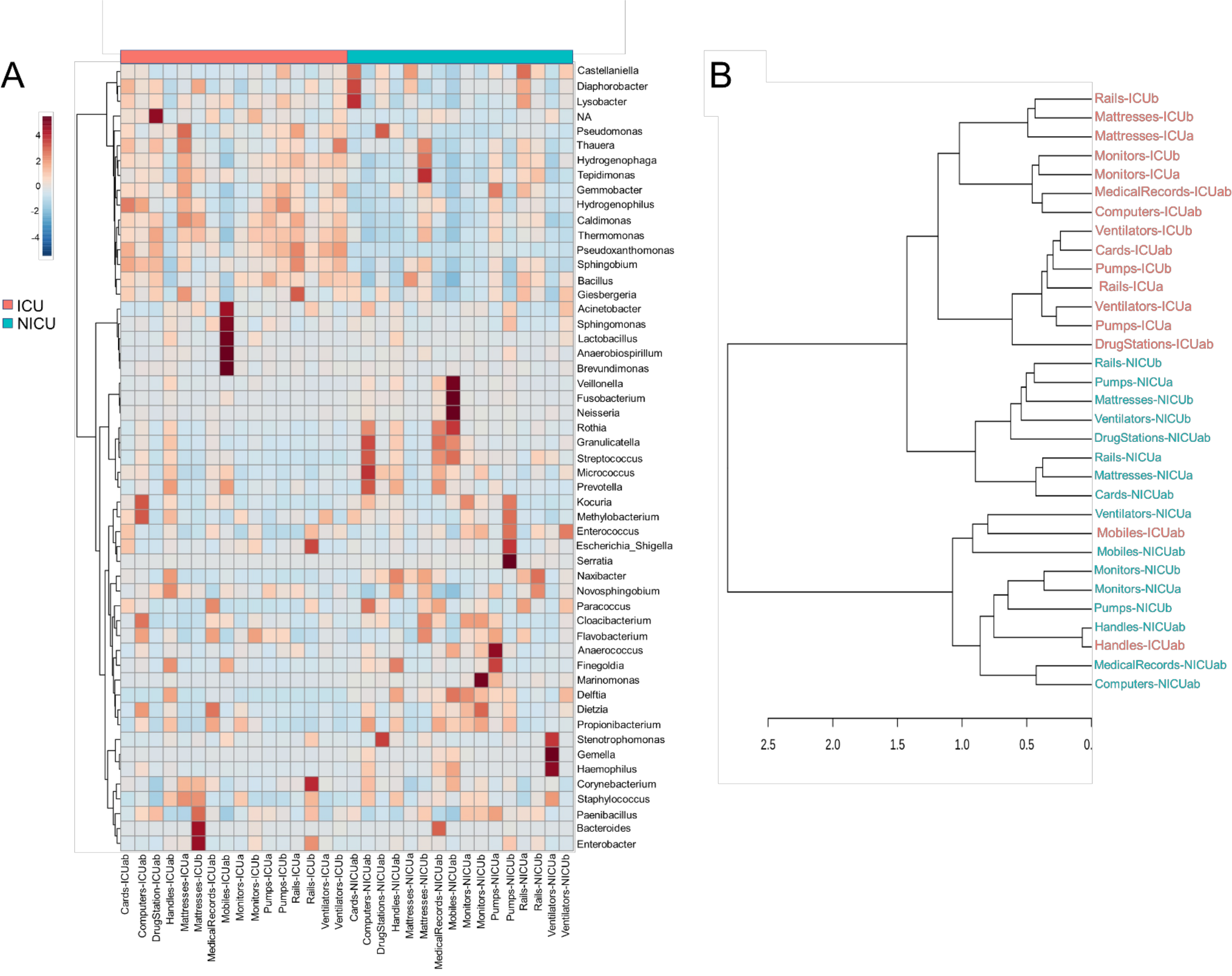
Clustering analysis of the ICU and NICU. (**A**) Heatmap, and hierarchical clustering of the main genera associated with ICU and NICU samples. The heatmap shows the relative abundance of the top 52 bacterial genera (rows) in each sample (columns). Hierarchical clustering is based on Ward Clustering algorithm and Euclidean Distance measure to generate the hierarchical tree. The color bar indicates the range of the relative abundance. (**B**) Dendrogram showing the similarities between ICU and NICU samples. The dendrogram was created using the Jaccard index as distance measure and Ward’s clustering algorithm.

Computers are indispensable in contemporaneous hospitals, and consequently, keyboard and mouse may be contaminated with dangerous pathogenic bacteria [29,30]. Here, we found potential opportunistic genera such as *Kocuria* (present at the skin and oral flora) and *Methylobacterium* in great abundance in ICU computers whereas NICU computers were enriched with *Rothia*, *Granulicatella*, *Streptococcus*, *Micrococcus*, and *Prevotella* (Fig. 3A). Another important, but generally neglected, potential vector of pathogens are the medical records (aka medical charts), especially those from (N)ICUs [31,32]. ICU medical records were enriched with *Dietzia* and *Flavobacterium*. NICU medical records were similar to NICU computers, except for being more abundant in *Bacteroides* (Fig. 3A). Moreover, fecal indicators were detected in a high proportion of NICU medical records (Fig. S3A). A hierarchical clustering analysis (Fig. 3B) based on the taxonomy of the ICU and NICU samples grouped them into two major clusters. Most of the samples from the same unit were clustered together indicating their similarity. Nonetheless, the microbiota community of ICU mobiles and handles were dispersed: mobiles-ICUab clustered closely with NICU ventilators (and mobiles), while ICU handles clustered with NICU handles group. These samples belonged to a cluster that revealed an almost absent *Bacillus* and higher frequency of *Streptococcus*, among other differences (Fig. 3A-B). Medical records were taxonomy similar to computers and also closer to monitors (Fig. 3B). Generally, for each unit, samples from surfaces frequently touched by HCW clustered together (Fig. S4A-B). These samples showed a higher abundance of skin-associated genera. The effects of these contamination sources for the patients were not part of this study. However, based on a vast literature, it is highly recommended to sensitize healthcare staff to sanitize mobiles, hands, computers and medical records (often neglected) to prevent cross-contamination within the hospital environment.

### Identification of ICU and NICU bacterial biomarkers

Across the ICU and NICU samples, different biogeographical patterns were observed for the different microbiotas. LEfSe analysis was performed to identify the distinguishing genera between ICU and NICU (Fig. 4A). LEfSe is a method that allows biomarker discovery most likely to explain differences between groups based on statistical significance, biological consistency and effect relevance [33]. In total, 25 genera were identified with LDA scores > 3.0. At the genus level, 11 specific biomarkers were present in NICU and 6 in ICU. All of them were both highly discriminatory and significantly different (p-value and FDR < 0.05) in term of abundances (Fig. 4B). The HAI-related genera *Delftia*, *Streptococcus*, *Haemophilus*, *Gemella*, *Serratia*, *Elizabethkingia*, *Leptotrichia*, *Clostridium*_*sensu*_*stricto*, *Chryseobacterium*, and *Vibrio* were biomarkers for NICU. Although most of these genera can be found in the respiratory tract, mouth, vagina, and intestinal tract of healthy adults, they present a high potential for nosocomial infection in neonates. Among these genera, there is a predominance of organisms with low oxygen tolerance (facultative anaerobes or obligate anaerobes). *Pseudomonas* was identified as a biomarker for ICU. It is well known that nosocomial infections caused by *Pseudomonas* are more often in ICUs than in other wards in the hospital [34]. Except for *Streptococcus* and *Leptotrichia*, all these HAI-related genera were found mainly in surfaces closer to the patients (boxes areas). Biomarkers could be used as indicators for the contamination status in a specific area in the hospital. Genera detected as biomarkers suggest that some bacteria can adapt extraordinarily within a particular environment.

**Figure 4.**
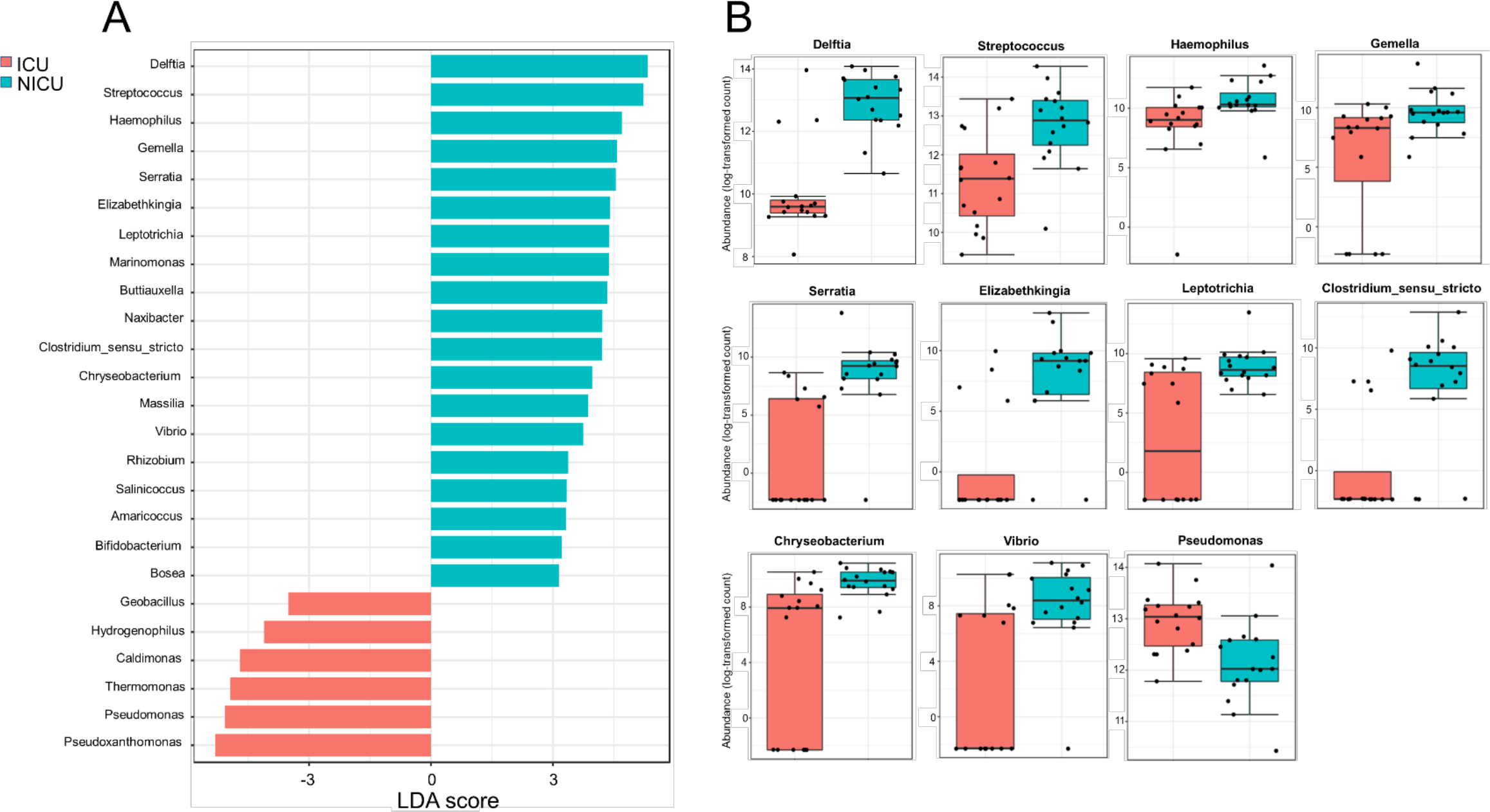
Significant differences between ICU and NICU. (**A**) Taxonomic biomarkers for ICU and NICU. Linear Discriminant Analysis (LDA) combined with Effect Size (LEfSe) indicate significant differences at the genus level that enable discrimination between the ICU and NICU samples (p < 0.05). Only those genera with log LDA score >3 are ultimately considered. (**B**) Boxplot of relative abundance (log scale) of the eleven HAI-related bacterial genera with significant differences between ICU (red, n=16) and NICU (cyan, n=16). The difference was calculated using Mann-Whitney/Kruskal-Wallis test (p-value and FDR < 0.05).

### ICU and NICU microbiotas have well defined community-level structures

Community-level relationships among the top 50 abundant bacterial genera were investigated through Pearson’s r correlation analysis (Fig. 5). Microbial interaction has an essential influence on antibiotic resistance and pathogenicity. In the ICU microbiome (Fig. 5A), five distinct clusters (i-v) were detected with significant positive correlations (co-occurrence). These clusters include potentially pathogenic genera such as (i) *Enterobacter*, *Staphylococcus*, *Corynebacterium*, and *Escherichia*_*Shigella*; (ii) Bacteria associated with outside environment (water, soil, and plants), among which *Pseudomonas*; (iii) *Stenotrophomonas*, *Acinetobacter*, *Sphingomonas*, and *Brevundimonas* (which can also cause co-infection with *Acinetobacter* spp.) [35]. (iv) *Enterococcus*, *Haemophilus*, *Kocuria*, *Dietzia*, *Gemella*, and *Neisseria*; (v) *Micrococcus*, *Fusobacterium*, *Prevotella*, *Delftia*, *Veillonella*, *Granulicatella*, *Rothia*, and *Streptococcus*. Except for *Pseudomonas*, the genera *Thermomonas*, *Bacillus*, and *Pseudoxanthomonas* showed negative correlations with all the five clusters cited above. In the NICU (Fig. 5B), we highlighted four (i-iv) clusters containing the following genera associated with nosocomial infections: (i) *Acinetobacter*, *Kocuria*, *Delftia*, and *Dietzia*; (ii) *Staphylococcus*, *Gemella*, and *Haemophilus*; (iii) *Fusobacterium*, *Neisseria*, *Corynebacterium*, *Rothia*, *Granulicatella*, and *Streptococcus*; (iv) *Enterobacter*, *Enterococcus*, *Sphingomonas*, *Escherichia*_*Shigella*, and *Serratia*. However, all these clusters revealed a strong negative correlation with *Bacillus*, *Sphingobium*, *Hydrogenophaga*, *Thauera*, *Thermomonas*, and *Gemmobacter*. It is important to note that most of these bacterial genera are known players in biofilms formation, including synergic multi-genera biofilms, on various hospital dry surfaces [36,37]. Biofilms matrix is a resistance mechanism that could stabilize a bacteria community in a selective environment such as (N)ICUs [38].

**Figure 5.**
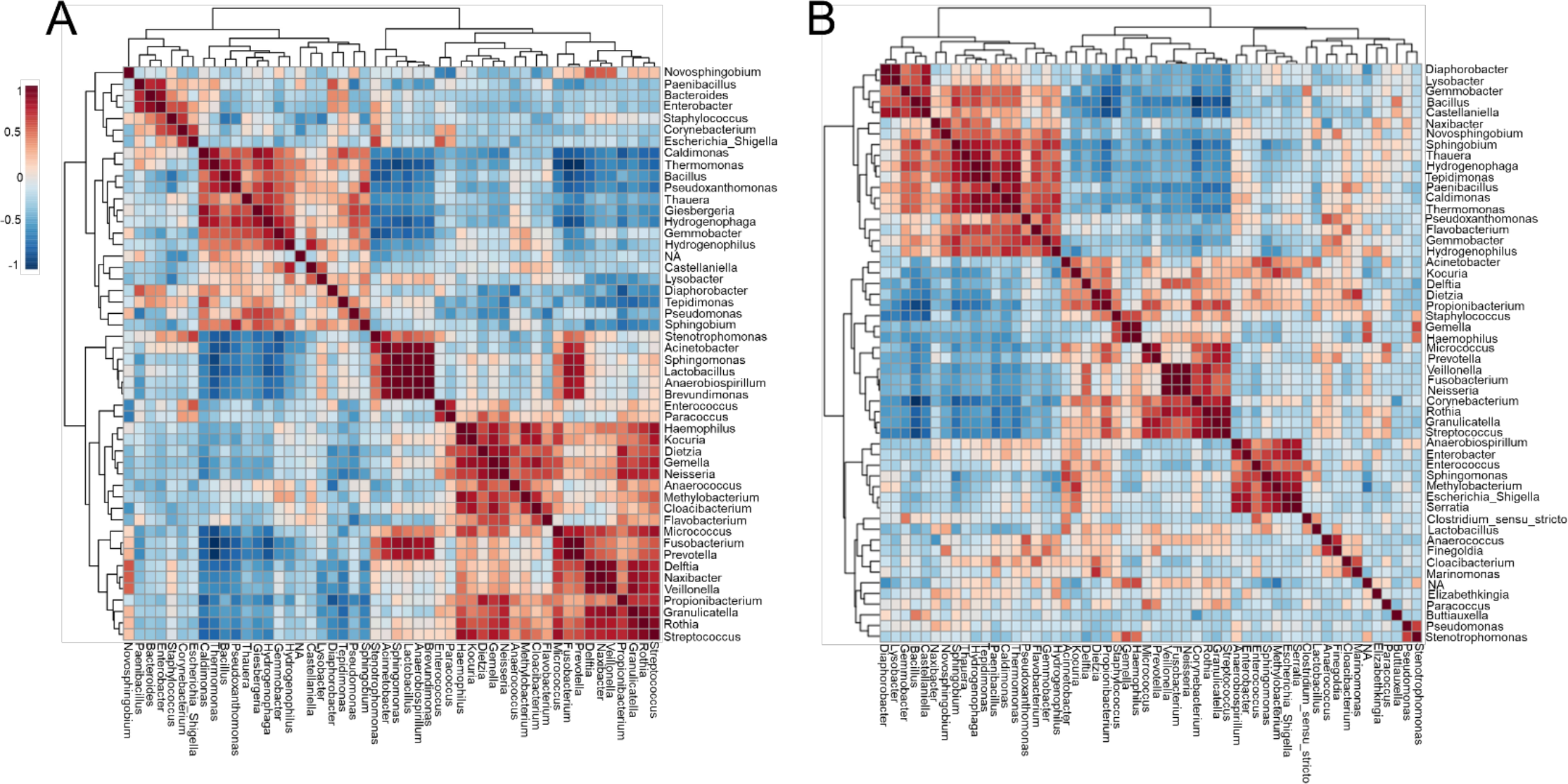
Co-occurrence and co-exclusion analysis of the bacterial genera. Heatmap showing Pearson’s r correlation coefficients among the top 50 abundant bacterial genera from the (**A**) ICU and (**B**) NICU. The correlation values ranged from −1.00 (blue) to 1.00 (red). Each square represents the Pearson’s r correlation coefficient between the genera of the column with that of the row. Self-self-correlations are identified in brown.

In order to verify whether the most prevalent potentially pathogenic genera identified in the ICU and NICU correlate with infected patients, 108 bacterial strains were isolated. Following standard cultivation, these strains were isolated from blood, bronchoalveolar lavage, peritoneal, cerebrospinal and ascitic fluids of hospitalized patients. All these isolates were identified, at the species level, by selective media, morphological features, and Vitek 2 rapid identification system and distributed among 12 genera. These strains comprised the genera *Klebsiella*, *Acinetobacter*, *Stenotrophomonas*, *Staphylococcus*, *Streptococcus*, *Pseudomonas*, *Enterobacter*, *Escherichia*, *Burkholderia*, *Cupriavidus*, *Morganella*, and *Ralstonia*. The most common culture-dependent isolates matched with the most abundant HAI-related genera found in the sequencing data (Fig. S5A-C). This correlation shows that potentially pathogenic organisms, even when found in abundance <10% in sequencing, may be predominant in hospital infections. The majority of the isolates obtained belonged to *Staphylococcus*, which was the second more abundant Gram-positive genus found in the sequencing. *Staphylococcus* already is described as one of the most common genera found in hospitals [39].

### Investigation of ICU microbial community profiling reveals substantial variation on the efficiency of the cleaning procedures

Cleaning procedures at ICUs are an important practice to prevent HAI-related bacteria spreading [39]. Although the protocols may vary between hospitals, concurrent cleaning procedures involved strict disinfection and sterilization of patient supplies and equipment during hospitalization. Here, the antimicrobial solution used for daily ICU cleaning contained the cationic polymer polyhexamethylene biguanide (PHMB). A recent model suggests that PHMB enter bacterial cells and condenses chromosomes, inhibiting cell division [40]. Thus, in order to investigate how concurrent cleaning affects the ICU microbiome, samples from surfaces near patients were sequencing and analyzed either before or immediately after cleaning. The microbial communities at genus level included sequences of 117 and 94 genera, for before and after cleaning respectively (Fig. 6A). Seven percent of the OTUs could not be classified to genera level (NA). These unclassified groups had higher relative abundance in cufflator-ICUab (35%). Samples after cleaning showed a slight but significant decrease in the diversity (Kruskal-Wallis test, p-value < 0.05) (Fig. 6B). However, noticeable variation was observed within the sample types (Fig. S6A). Beta diversity analysis revealed distinct, but overlapping, profile (R= 0.091961; p-value < 0.05) (Fig. 6C). Most of the samples from ICU ward-A after cleaning clustered separately from the rest of the surfaces. Quite remarkably, these differences in diversity after cleaning reveal that the procedure did not have the same effect on all surfaces. Although it is known that different microbiomes may exert different effects on cleaning [37], this was not the case, since no significant difference between room A and B was observed prior to cleaning. Therefore, differences in the effect of cleanliness on diversity could be explained, in part, by a lack of standardization in the protocol.

**Figure 6.**
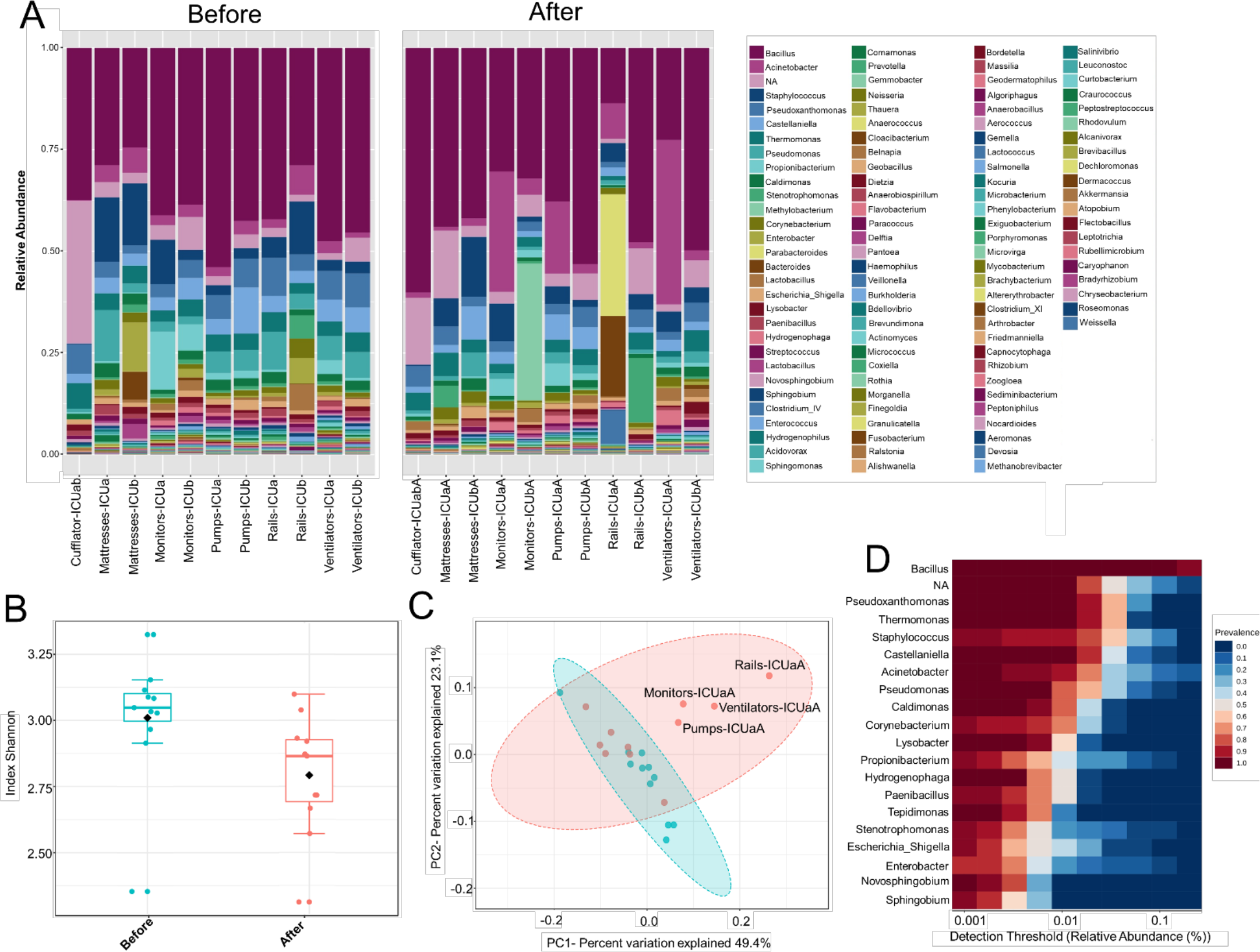
ICU bacteria microbiota profile before and after cleaning. (**A**) Relative bacterial abundance at the genus level. Sequencing results are showed for each sample surface clustered using Usearch algorithm with a 97% cutoff. Only genera with abundance > 1.0% were plotted. (**B**) Alpha diversity at OTU level, before (red, n=11) and after cleaning (cyan, n=11) calculated using Shannon index (Kruskal-Wallis test, p-value < 0.05). (**C**) PCoA plot based on Jensen-Shannon distances between bacterial communities associated with cleaning (ANOSIM, R= 0.091961; p-value = 0.039). Samples are shown as single dots. Divergence at OTU level was computed on Total sum scaling–normalized (TSS-normalized) datasets. (**D**) Core microbiome analysis based on relative abundance and sample prevalence of bacterial genus before and after cleaning.

The samples either before or after cleaning were inhabited by high relative abundances (~65%) of *Bacillus*, *Pseudoxanthomonas*, *Thermomonas*, *Staphylococcus*, *Castellaniella*, and *Acinetobacter*. Core microbiome analysis showed that 19 genera were shared in 80% of all samples (before and after) at the minimum detection threshold of 0.001% relative abundance (Fig. 6D). Most notably, the most abundant genera were also clearly most prevalent in the core microbiome before and after cleaning. Gram-positive bacteria were found in higher abundance (before – 53%; after – 51%, respectively), showing 45 different genera before and 30 after cleaning (33% less). Furthermore, Gram-negative bacteria revealed higher diversity, with 72 genera before and 64 after cleaning (11% less). Most of the genera absent after cleaning showed very low abundance (<0.05%) before cleaning. The HAI-related organism *Chryseobacterium*, and *Clostridium*_*XI* are among the genera absent (or extremely low) after cleaning. Besides these absent genera, using the statistical parameters p-value and FDR < 0.05, no other analyzed genera showed a significant difference between the average abundance calculated for all samples before and after cleaning. However, the HAI-related genera *Comamonas*, *Pseudomonas*, *Enterobacter*, *Kocuria*, *Ralstonia*, and *Delfitia* showed a decrease, while *Leptotrichia*, *Streptococcus*, and *Acinetobacter* presented an increase on average abundance ≥two-fold after cleaning (Fig. S7A). Curiously, cleaning efficiency was notably variable among the samples (Fig. S7B). Previous studies have shown that even with strict cleaning procedures, HAI-related genera, such as *Staphylococcus*, *Klebsiella*, *Acinetobacter*, *Pseudomonas*, *Enterococcus*, *Escherichia*, and *Enterobacter*, are generally found on the surface of the ICU devices [41–45]. To examine more deeply the cleaning effect among the samples, a heatmap of the top 45 genera is illustrated in Fig. 7A. The cleaning efficiency was not the same through the samples and wards. Some genera showed a tendency to decrease after cleansing, such as *Enterococcus*, *Enterobacter*, *Staphylococcus*, *Burkholderia*, *Comamonas*, *Pseudomonas*, and *Delftia*. However, others increased in one ward and dropped in the other, such as *Corynebacterium* and *Acinetobacter* (increased for ward-A and decreased for ward-B) or *Prevotella* and *Novosphingobium* (decrease for ward-A and increase for ward-B).

**Figure 7.**
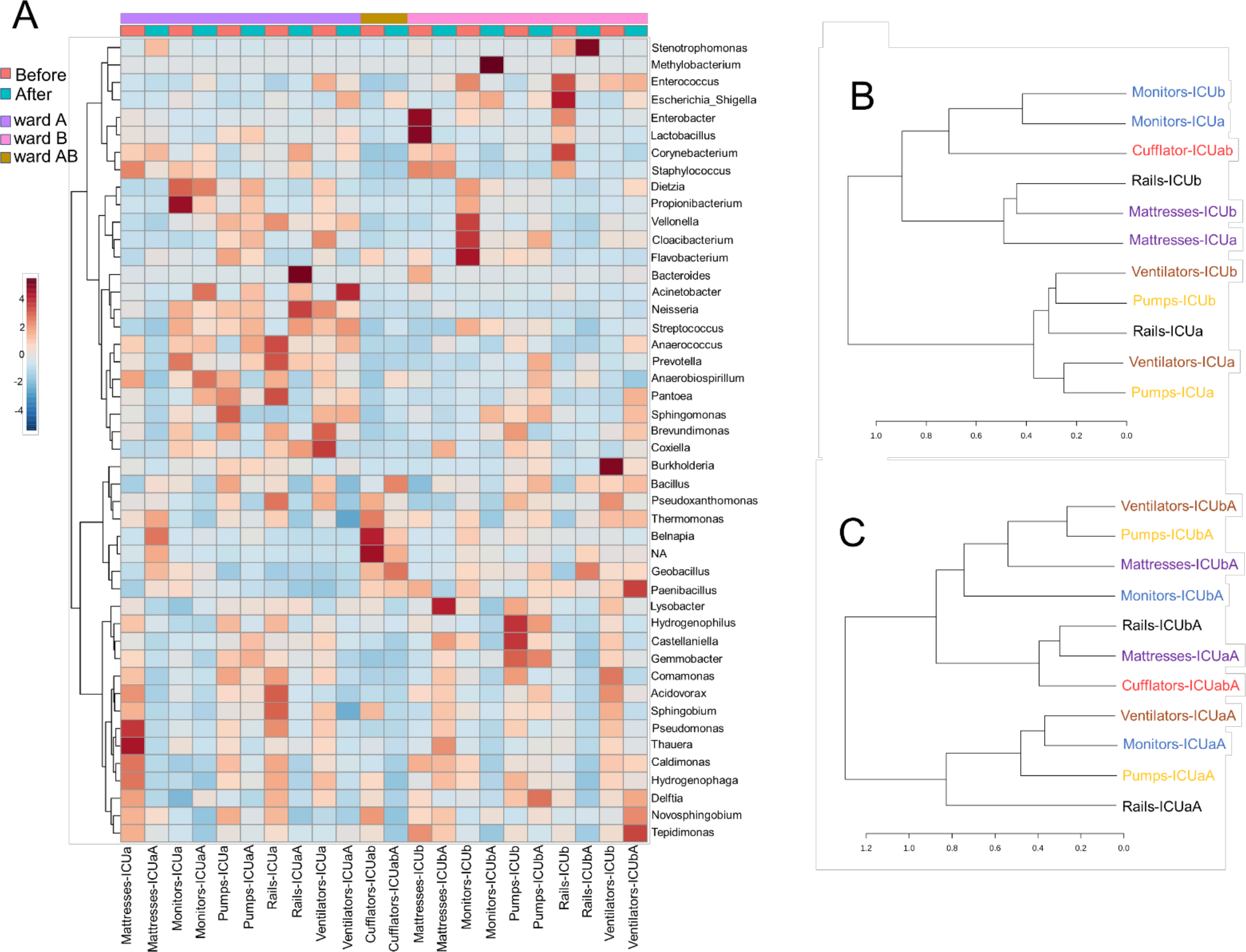
Clustering analysis of the ICU samples before and after cleaning. **(A)** Heatmap of the main genera associated with ICU samples before and after cleaning. The heatmap shows the relative abundance of top 45 bacterial genera (rows) in each sample (columns). The color bar indicates the range of the relative abundance. Dendrogram showing the similarities between samples **(B)** before and **(C)** after cleaning. The dendrogram was created using the Jaccard index as distance measure and Ward’s clustering algorithm.

Moreover, there were genera that revealed an tremendous increasing after cleaning in some specific surfaces, such as *Stenotrophomonas* (mattresses-ICUaA and rails-ICUbA), *Methylobacterium* (monitors-ICUbA), *Bacteroides*, *Neisseria*, and *Streptococcus* (rails-ICUaA), *Acinetobacter* and *Escherichia* (ventilators-ICUaA), *Dietzia* (monitors-ICUaA), *Delftia* (pumps-ICUbA), *Novosphingobium* and *Tepidimonas* (ventilators-ICUbA). Fecal indicators were detected in higher abundance after cleaning on bed rails (mainly on Rails-ICUaA) (Fig. S3A). These results reveal that cleaning was inconsistent and, in some cases, increased the load of specific genera. Previous studies have shown that hands are one of the primary vectors of HAI-related bacterial cross-contamination [46,47], mainly because of the variable compliance on hands hygiene and gloves changing after touching surfaces near to the patients [48]. Besides, disinfectant solutions and wipes used for hospital cleaning also can be a vital source of pathogen transfer and inconsistency in surfaces cleaning, even when standard protocols are followed [49]. Furthermore, other factors to be considered is the low efficiency of PHMB-based products in relation to contaminations by wound secretions or urine containing a massive load of bacteria [50], and a possible discrepancy in the cleaning procedure performed by different nurses. Based on hierarchical clustering analysis, before cleaning (Fig. 7B) most of the samples with the same functionality, but from different wards, were clustered together indicating their similarity. Nonetheless, the microbiota community after cleaning (Fig. 7C) revealed a higher dispersion among the samples. We speculate that cleaning could be a way of spreading colonizing genera from one surface to another, but that over time there may be a reestablishment of the microbial community related to a specific sample.

### Cleaning procedures generates substantial rearrangements in the community-level structures

To investigate the changes in the microbial community structure before and after cleaning the correlation coefficients among the top 50 genera was analyzed (Fig. 8A-B). For the microbiome before cleaning, four distinct clusters (i-iv) were detected with significant positive co-occurrence (Fig. 8A). These clusters include potentially pathogenic genera such as (i) *Enterococcus*, *Escherichia_Shigella*, *Stenotrophomonas*, *Enterobacter*, *Staphylococcus*, *Acinetobacter*, and *Corynebacterium*; (ii) *Dietzia*, *Streptococcus*, and *Veillonella*; (iii) *Sphingomonas*, *Neisseria*, and *Methylobacterium*; (iv) *Burkholderia*, *Pseudomonas*, *Ralstonia*, and *Comamonas*. The environmental genus *Belnapia* showed negative correlations with all the genera cited above.

**Figure 8.**
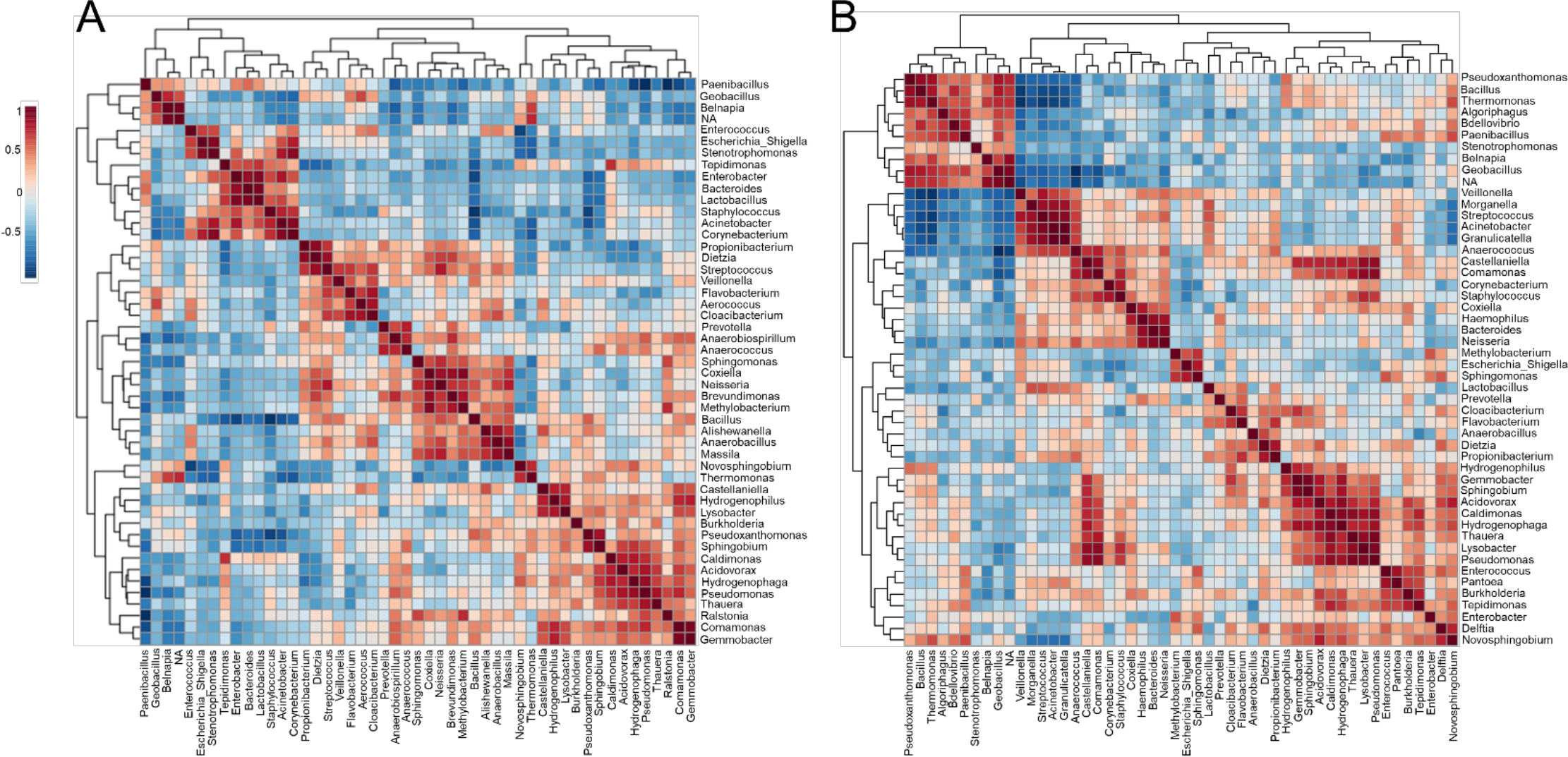
Co-occurrence and co-exclusion analysis of the bacterial genera. Heatmap showing Pearson’s r correlation coefficients among the top 50 abundant bacterial genera from the (**A**) ICU before and (**B**) ICU after cleaning. The correlation values ranged from −1.00 (blue) to 1.00 (red). Each square represents the Pearson’s r correlation coefficient between the genera of the column with that of the row. Self-self-correlations are identified in brown.

After cleaning, six clusters (i-vi) are presented (Fig. 8B) containing highlighted genera associated with nosocomial infections: (i) *Stenotrophomonas* and other environmental genera; (ii) *Veillonella*, *Morganella*, *Streptococcus*, *Acinetobacter*, *Granulicatella*, *Comamonas*, *Corynebacterium*, *Staphylococcus*, *Haemophilus*, and *Neisseria*; (iii) *Methylobacterium*, *Escherichia*, and *Sphingomonas*. (iv) *Prevotella* and other genera related to low oxygen tolerance or vaginal microbiome [51]; (v) *Pseudomonas*, *Enterococcus*, *Pantoea*, and *Burkholderia*; (vi) *Enterobacter*, *Delftia*, and *Novosphingobium*. However, most of these HAI-related genera revealed a strong negative correlation with *Pseudoxanthomonas* (except *Delftia* and *Novosphingobium*). The correlation data showed a predominance of Proteobacteria among most of the clusters. Proteobacteria are predominant in the skin of the forearm [52] and are highly associated with biofilms formation on the surface of devices used on ICUs [36]. Several genera relationships were quite stable to disinfection stress because it was found clustered both before and after cleaning. In all the clusters were found genera associated with species able to form biofilms. Genera associated with xenobiotic metabolism were found among the clusters i-iv, and ii-v before and after cleaning, respectively [53,54]. After cleaning it was noticed a redistribution of some genera in new clusters. For example, a more extensive cluster involving ten HAI-related genera (ii) was formed after cleaning, this cluster included a mixture of several genera found in clusters i-iv before cleaning. Although this cluster analysis is useful to visualize the dynamics of microbiotas with the cleaning efficiency, further studies will be required to understand the exact changes in the microbe-microbe interactions underlying the differences observed across time.

## Conclusions

The relevance of spatial composition of the microbial communities within a hospital is unclear. To our knowledge, this is the first study using deep sequencing of inanimate surfaces samples to develop a spatial assessment of the microbial community in ICU and NICU wards within the same hospital. In this comprehensive study, we observed a peculiar spatial structure between ICU and NICU microbiota in one of the largest hospitals in Brazil. The data revealed that among the samples analyzed, NICU presents higher biodiversity than in the ICU. Genera considered “survival specialists” are among the most persistent and abundant in both units. Areas closest to the patient hold more specific microbiota, distinguishing one unit from other. Most of the genera found in both units are present on the healthy human microbiome, suggesting that the most likely vectors of contamination are hospital staff and patients. Most of these genera can also be associated with nosocomial infection, especially for patients in (N)ICU. Devices commonly used, but generally neglected, such as mobile phones, computers, and medical charts are enriched with HAI-related genera (e.g., *Acinetobacter*, *Fusobacterium*, *Kocuria*, *Rothia*, and *Dietzia*). For the samples analyzed in the present study, some facultative anaerobes or obligate anaerobes genera were classified as biomarkers for the NICU (e.g., *Serratia* and *Clostridium*), whereas *Pseudomonas* as a biomarker for ICU. Correlation analyses revealed a distinct pattern of microbe-microbe interactions for each unit, including several bacteria able to form multi-genera biofilms. Cultivation-dependent results showed a positive correlation between the most abundant HAI-related genera identified by sequencing with infections found in the hospital. According, our data showed similarity with previous studies and can help to define soon what constitutes a "typical" microbiome in the ICU and NICU environments. The ability to identify HAI-related genera that are spatially concentrated in a hospital ward may influence the future use of improved tools and protocols for infection control.

Furthermore, we evaluated the effect of concurrent cleaning over the ICU bacterial community. Cleaning showed a slight decrease in diversity. Several genera were quite stable to disinfection, suggesting being well-adapted to the ICU environment. In general, the cleaning procedure was inconsistent. Potential influencing factors from the unsatisfactory cleaning include low efficiency of the biocide used, bacteria well-adapted to daily cleaning, disinfectant solutions and wipes contaminated, and variable compliance on hands hygiene and cleaning procedure. Therefore, this type of analysis can be used for designing better strategies for cleaning procedures. In conclusion, we demonstrate here how NGS could be used for monitoring potential contamination sources in (N)ICU units and to evaluate existent decontamination protocols stablished in these unities. We highlight that similar approaches, while still very costly, could be implemented for periodic monitoring of microbial profiles in clinical hospitals to help reducing potential secondary infections.

## MATERIALS AND METHODS

### Sample collection and DNA extraction

A total of 158 samples were collected from the ICU and NICU at The Medical School Clinics Hospital (Ribeirão Preto, Brazil) by a single investigator from September to October 2018. The intensive care units contained two wards with four beds each, where critically ill patients from all medical specialties are treated. Samples from NICU were collected only before the concurrent cleaning, while from ICU samples were collected either before or immediately after cleaning. During sampling, all employees and devices of the ICU/NICU were in full operation. Boxes with patients lying down were swabbed on the surfaces of mattress, bed rail, monitors, infusion pumps, ventilator, and cufflator (when present). In common areas of the ICU/NICU, computer keyboard and mouse, doors handle, hospital cards, medical records, drug station, and nurse’s mobiles were also swabbed. All sampling locations and their characteristics are given in Fig.1 and Table 1. The following code was used to name the samples: Samples-Unit (ICU or NICU) ward (a, b or ab) A (after cleaning), e.g., Monitors-ICUaA Samples were collected using sterile swabs (Absorve^®^, Jiangsu, China) premoistened with sterile Amies media [55]. The swabs were streaked across a 400-cm^2^ area in four different directions with firm movements for 2 minutes; swabs were rotated to ensure full contact of all parts of the swab tip and the surface. After a surface was sampled, the swab was immediately placed into sterile 15-ml Falcon tubes containing 1 mL of sterile Amies media and stored in a 4°C cooler until returning to the laboratory. In the laboratory, due to extremely low biomass, samples from a similar source and the same ward were pooled together –, e.g., four monitors from NICU ward A is a pool, and four monitors from NICU ward B another pool– generating 43 pooled samples. Then, the samples were concentrated to 500 μL by centrifugation (10000 g / 20min), and DNA was extracted using the MoBio Powersoil DNA isolation kit, then stored in a −80°C freezer until further processing.

### Concurrent cleaning procedures in the ICU

At the beginning of each 24-h shift, a registered nurse washed his or her hands, put on nonsterile gloves, and wiped Boxes surfaces (mattress, bed rail, computer touch screens, monitors, infusion pumps, ventilator, and cufflator) with 1% polyhexamethylene biguanide (PHMB) solution on a soft wipe.

### Sequencing and diversity analysis

The DNA concentrations were measured fluorometrically (Qubit® 3.0, kit Qubit® dsDNA Broad Range Assay Kit, Life Technologies, Carlsbad, CA, USA). DNA integrity was determined by agarose gel electrophoresis using a 0.8% (w/v) gel, and subsequent staining with SYBR Safe DNA Gel Stains (Invitrogen, Carlsbad, CA, USA). A PCR was employed to amplify the V4 regions of the 16S ribosomal RNA gene 16S rRNA for bacteria [17]. Each PCR reaction mixture contained 20 ng of metagenomic DNA, 10 μM of each forward and reverse primers, 1.25 mM of magnesium chloride, 200 μM of dNTP mix (Invitrogen, Carlsbad, CA, USA), 1.0 U Platinum Taq DNA polymerase high fidelity (Invitrogen, Carlsbad, CA, USA), high fidelity PCR buffer [1X], and milli-Q water. Reactions were held at 95 °C for 3 min, with amplification proceeding for 30 cycles at 95 °C for 30 s, 53.8 °C for 30 s, and 72 °C for 45 s; a final extension of 10 min at 72 °C was added to ensure complete amplification. The expected fragment length of PCR products was verified by agarose gel (1%) electrophoresis, and the amplicon size was estimated by comparison with a 1 kb plus DNA ladder (1 kb plus DNA ladder, Invitrogen, Carlsbad, CA, USA). The PCR fragments were purified using the Zymoclean™ Gel DNA Recovery kit following the manufacturer’s instructions. Sequencing was performed using the Miseq Reagent kit v3 2 × 300 bp.

All sequences data were processed, removing adapters using Scythe 0.991 (https://github.com/vsbuffalo/scythe) and Cutadapt 1.7.1 [56]. Sequence trimming was carried out by selecting sequences over 200 bp in length with an average quality score higher than 20 based on Phred quality, and duplicate reads were removed using the Prinseq program [57]. The QIIME software package version 1.9.1 was used to filter reads and determine Operational Taxonomic Units (OTUs) as described in Caporaso et al. (2010). The Usearch algorithm was used to cluster the reads OTUs with a 97% cutoff, and to assign taxonomy using the Ribosomal Database Project (RDPII) version 10 [58]. Bacterial sequences were de-noised, and suspected chimeras were removed using the OTU pipe function within QIIME. Sequence data were summarized at the phylum, class, and family levels; Also, Alpha_diversity.py in QIIME was used to calculate ACE, Chao1, Shannon, and Simpson indices. Principal coordinate analyses (PCoA) were conducted to evaluate differences in community structure among experimental groups (β-diversity).

For further statistical analysis and visualization, OTU table with taxa in plain format and metadata file were uploaded to the MicrobiomeAnalyst tool (available at http://www.microbiomeanalyst.ca) [59]. Shallow abundant features were filtered using options; minimum count 4, low-count filter based on 20% prevalence in samples. For comparative analyses, a low variance filter was applied based on Inter-quantile range and removing the 10% lowest features. Data were rarefied to the minimum library size and normalized using total sum scaling (TSS) before any statistical comparisons [60].

## Availability of supporting data

The nucleotide sequences obtained in the present study have been deposited in the GenBank database under the Accession number PRJNA541082.

## Ethics approval and consent to participate

No specific permissions or ethics approval were required for this study with inanimate surfaces.

## Author Contributions

LFR, RSR, and MEG conceived of the project. LFR, LFCR, MGM, and GGG organized the sample collections. LFR conducted the nucleic acid extractions. EML and LTK conducted the MiSeq library preparations and provided the bioinformatics support, and LFR contributed to the data analysis. LFR, RSR, and MEG wrote the final manuscript. All authors have read and approved the manuscript.

## Supplementary material

**Figure S1.**
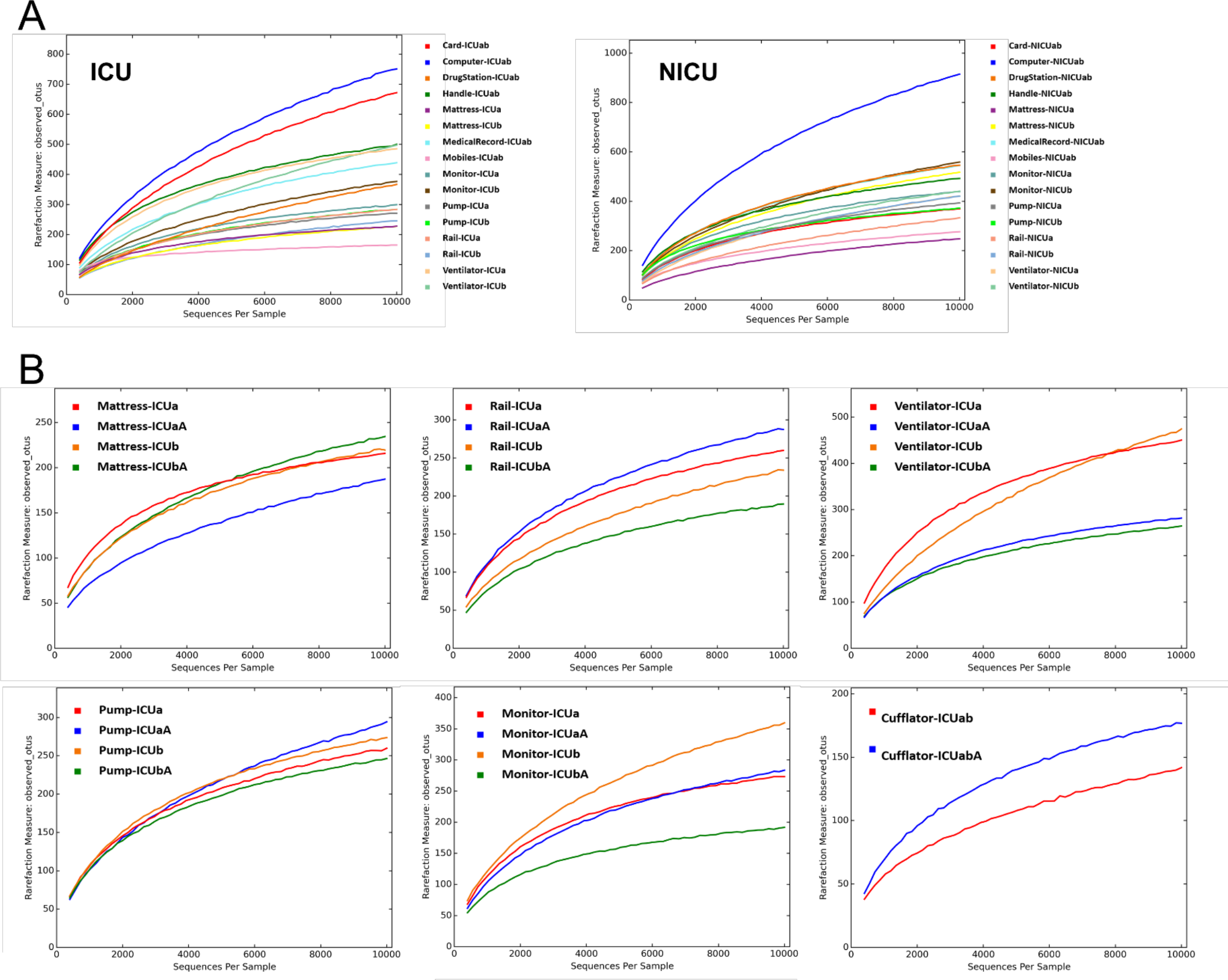
Rarefaction curve showing the relationship between the sequencing per sample and the number of OTUs that these reads represent. **(A)** ICU and NICU, and **(B)** ICU before and after cleaning. Sequences were rarefied with 33.708 read counts per sample.

**Figure S2.**
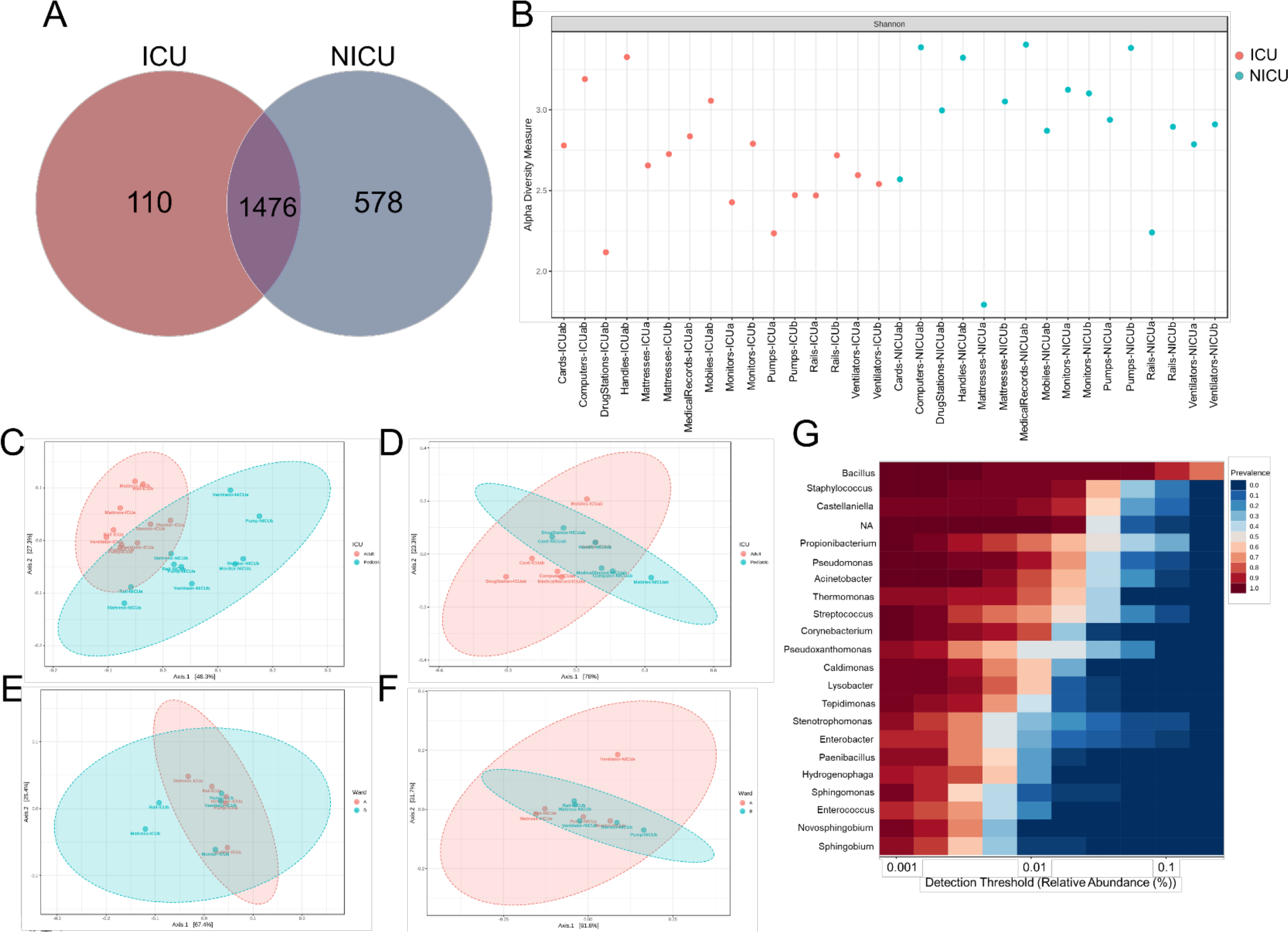
(**A**) Venn diagram showing shared and unique OTUs. (**B**) Alpha diversity at the OTU level for each sample at ICU (red) and NICU (cyan) calculated using Shannon index (Kruskal-Wallis test, p-value < 0.05). PCoA plot based on Jensen-Shannon distances between bacterial communities associated with (**C**) ICU and NICU boxes areas (ANOSIM, R= 0.50756; p-value < 0.001); (**D**) ICU and NICU common areas (ANOSIM, R= 0.14074; p-value = 0.116); (**E**) ICU wards (ANOSIM, R= 0.124; p-value = 0.177); (**F**) NICU wards (ANOSIM, R= −0.02; p-value = 0.52). Samples are shown as single dots. (**G**) Core microbiome analysis based on relative abundance and sample prevalence of bacterial genus in ICU and NICU. Divergence at OTU level was computed on Total sum scaling–normalized (TSS-normalized) datasets.

**Figure S3.**
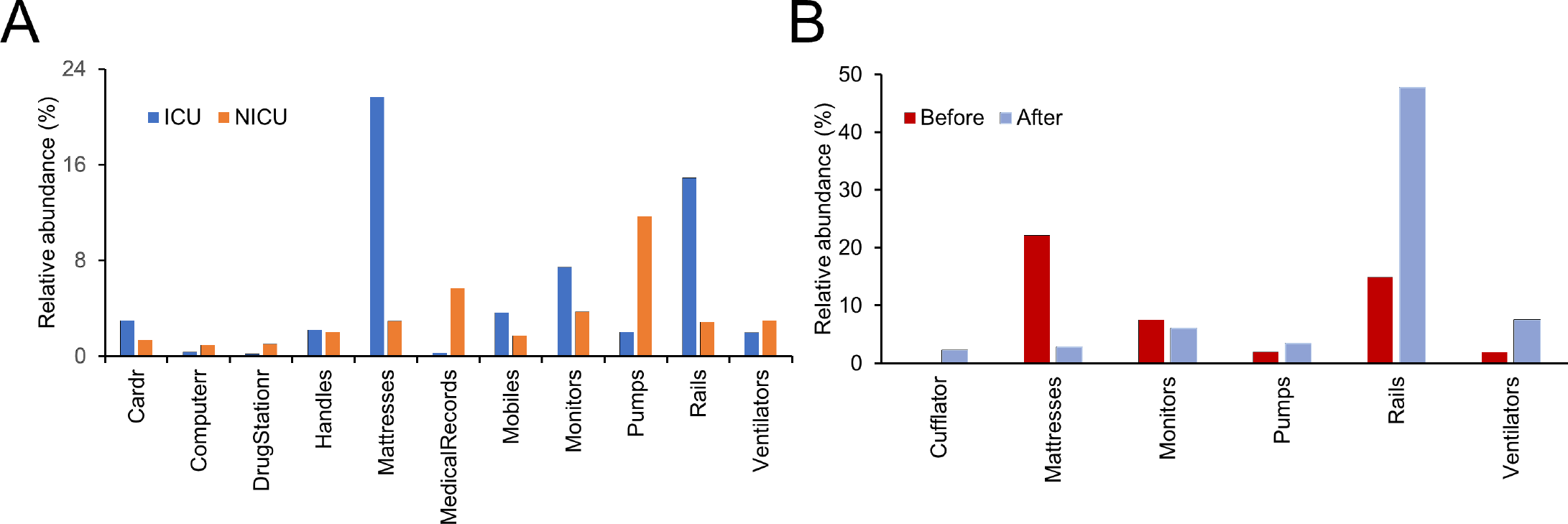
The abundance of fecal indicators in **(A)** ICU/NICU and **(B)** ICU before/after cleaning.

**Figure S4.**
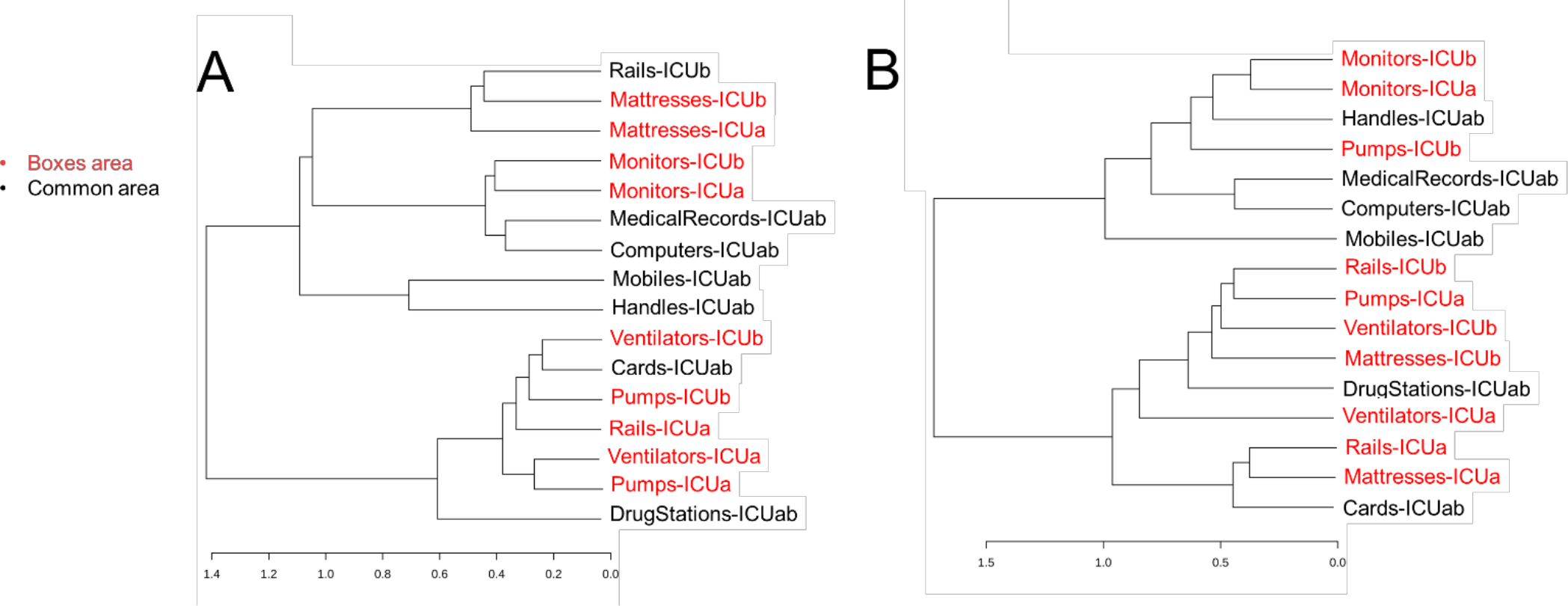
Dendrogram showing the similarities between samples. **(A)** ICU; **(B)** NICU. The dendrogram was created using the Jaccard index as distance measure and Ward’s clustering algorithm.

**Figure S5.**
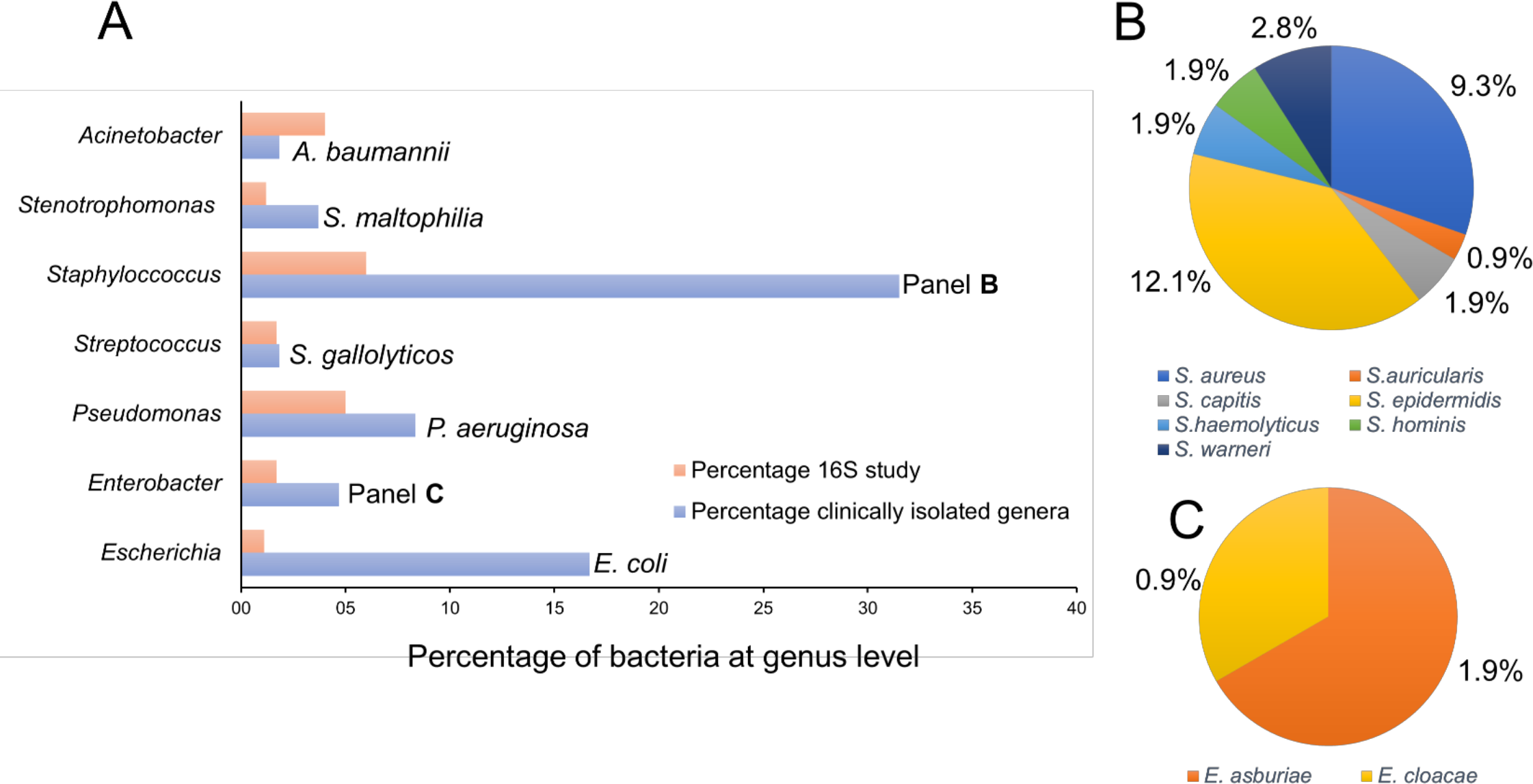
Percentage of identified bacteria at the genus level from the 16S analysis and isolated bacteria from clinical samples. (**A**) A total of 108 bacterial strains (gathered in 12 different genera) isolated from blood, bronchoalveolar lavage, peritoneal, cerebrospinal and ascitic fluids of hospitalized patients in the ICU were evaluated. In the graph are represented only the seven most abundant genera from the 16S amplicon study (considering up to 4% of all quality sequences). Data at the level of species are presented just for the bacteria isolated from clinical samples. Percentage of species belonging to the genera *Staphylococcus* (**B**) and *Enterobacter* (**C**), concerning the 108 bacterial strains isolated. For the other genera, single species were identified among biological samples, as presented in panel A.

**Figure S6.**
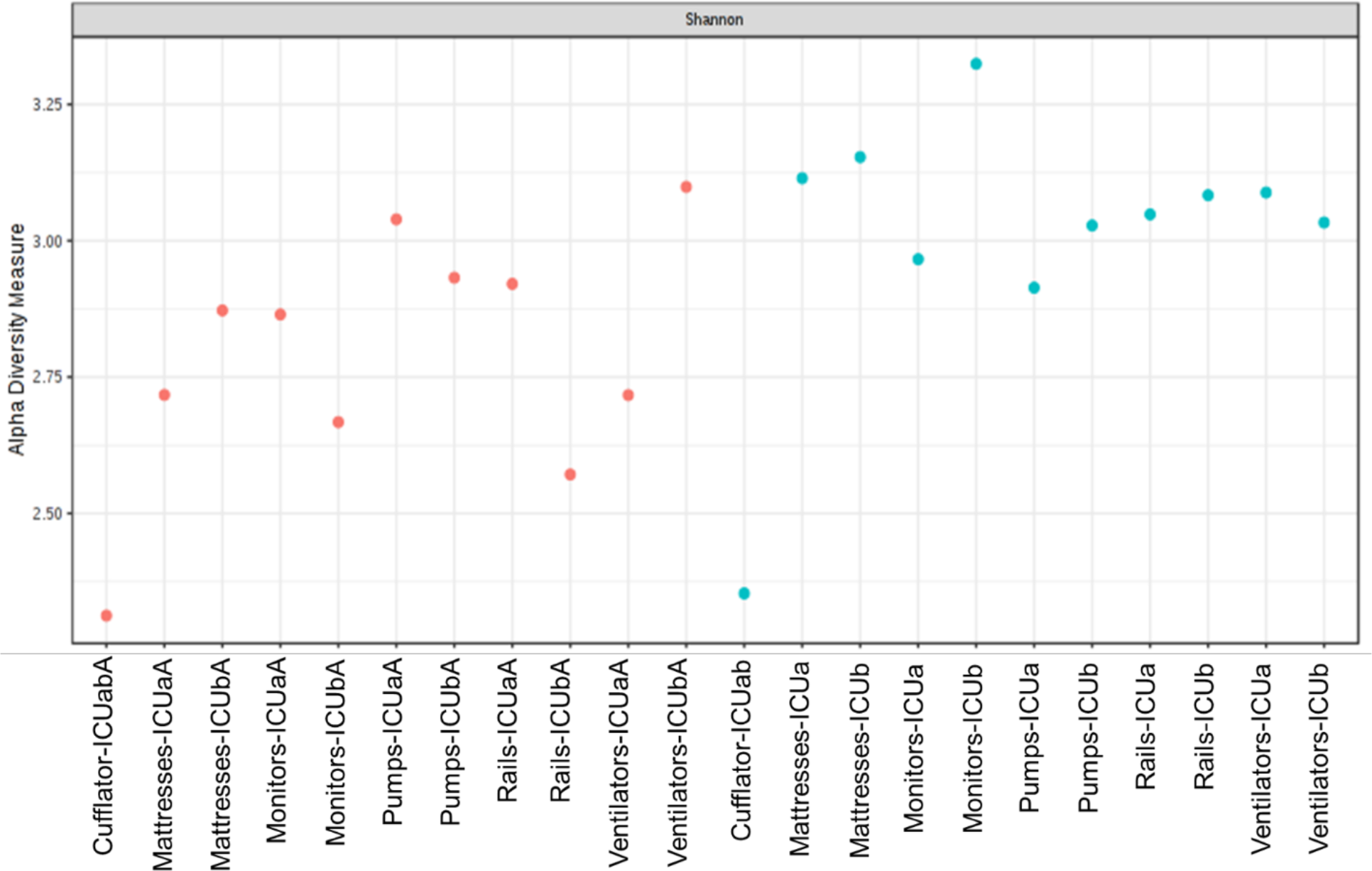
Alpha diversity at OTU level for each sample at ICU before (red) and after cleaning (cyan) calculated using Shannon index (Kruskal-Wallis test, p-value < 0.05).

**Figure S7.**
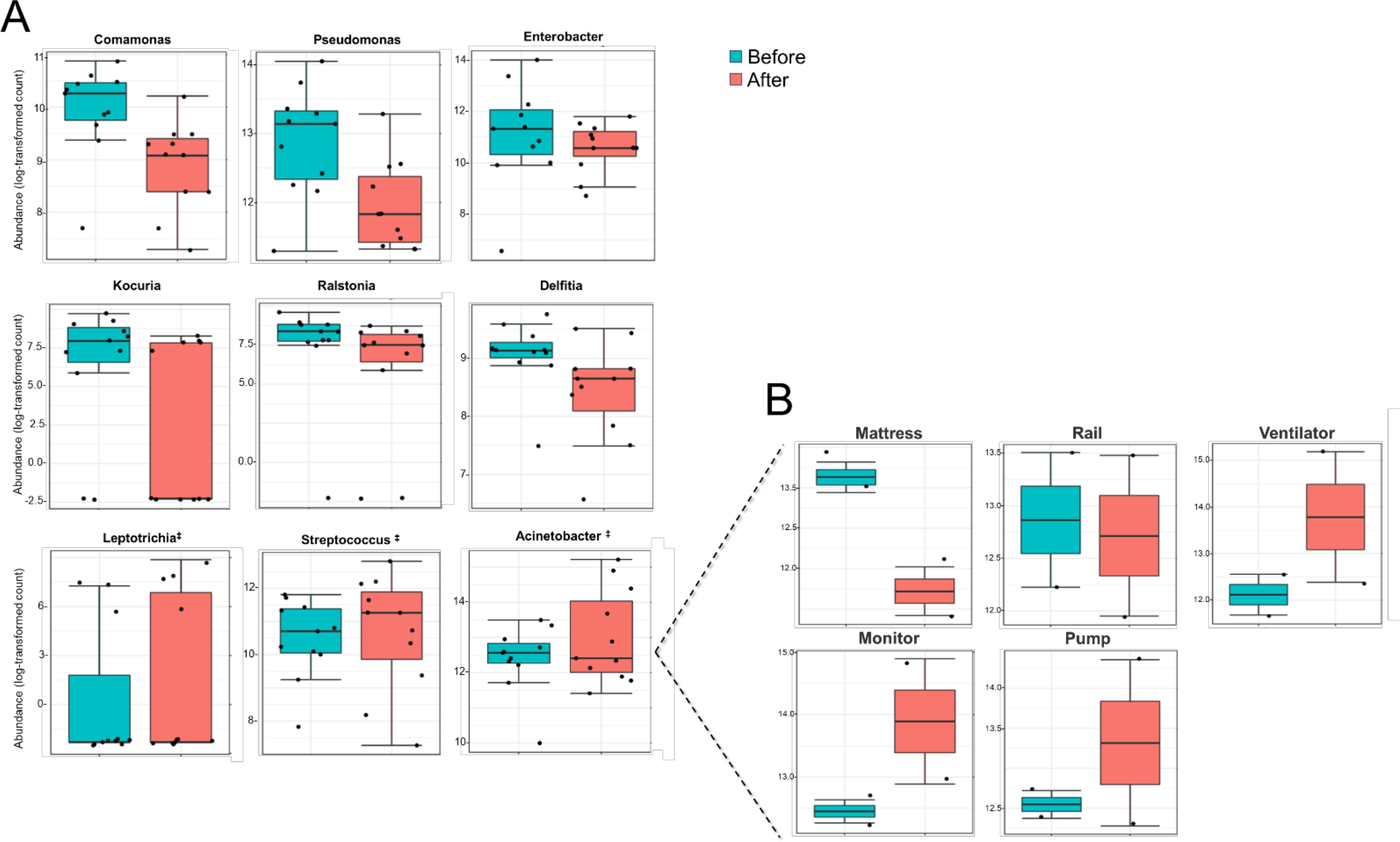
(**A**) Boxplot of relative abundance (log scale) of the genera HAI-related before (cyan) and after (red) cleaning. The difference was calculated using Mann-Whitney/Kruskal-Wallis test (FDR ≥0.3). ^‡^Genera with higher abundance after cleaning. (**B**) Differential relative abundance for *Acinetobacter* across all the ICU samples.

## Supplementary text

### Oxygen tolerance

Most of the samples contained a mixture of organisms with various degrees of oxygen tolerance. The number of strictly aerobic genera were highly represented (50%) followed by facultative anaerobe (36%) and obligatory anaerobic bacteria (14%) for both units. Infections caused by anaerobic bacteria are often underestimated, due to the difficulty to isolate and identify these microorganisms. The use of unspecific therapy against these infections may cause clinical failures [61]. Most abundant anaerobic organisms in ICU were, on decrescent order, *Propionibacterium*, *Bacteroides*, and *Prevotella*, whereas for NICU were *Propionibacterium*, *Prevotella*, and *Veillonella*. *Propionibacterium* is a human skin-associated genus [62], while *Prevotella* and *Veillonella* are part of the healthy microflora in the oral cavity and vaginal [51,63]. However, many species of *Prevotella*, and *Veillonella* genera are pathogens that cause oral or respiratory diseases [64], as well as meningitis [64]. *Veillonella* has also been involved in prosthetic cardiac valve or joint infections [65,66] and fatal sepsis [67]. *Bacteroides* species are usually part of the gastrointestinal microbiota [68], and they make a significant portion of the fecal bacterial population [69]. Among all anaerobic bacteria, *Bacteroides*, *Prevotella*, and *Veillonella* are the most frequently isolated in clinical samples of infection [70].

### Gram-positive bacteria

Gram-positive bacteria were found in higher abundance in both units (Gram-positive and Gram-negative at ICU – 49% and 46%; NICU – 52% and 44.5%, respectively; ~5% were Gram-variable). Nonetheless, in terms of the number of genera, Gram-negative bacteria were predominant in both ICU (70%) and NICU (66%). Over the last decades, the focus of infection control centers was targeting Gram-positive pathogens due to their high rate of morbidity and mortality [71]. However, the incidence of infections in UTIs caused by Gram-negative bacteria has been rising alarmingly, requiring a better understanding of hospital microbiomes [72–74].

The five more abundant Gram-positive genera found in both ICU and NICU samples were, in order of decreasing abundance, *Bacillus*, *Staphylococcus*, *Propionibacterium*, *Streptococcus*, and *Corynebacterium*. Core microbiome analysis was performed combining samples from both ICU and NICU. In total, 21 genera were shared in 80% of all samples at the minimum detection threshold of 0.001% relative abundance (Fig. S2G). The most abundant Gram-positive genera were also among the top 10 more prevalent. Most notably, *Bacillus* was the most prevalent genus in the core microbiome for both care units. *Bacillus* was also most abundant in ICU (36%) and NICU (26%) (Fig. 2A), mainly in the boxes area. To examine more deeply the bacterial community variations among the samples, a heatmap of the top 52 genera is illustrated in Fig. 3A. Accordingly, the ICU pumps and NICU mattresses contained the highest abundance of the total reads (~6%) (Fig. 3A). The identification of the *Bacillus* genus in hospital samples is often considered clinically safe since it is ubiquitous in the environment. However, recently outbreaks of severe and lethal *Bacillus* infections have been widely reported, especially diseases related with *Bacillus cereus* at NICUs [75,76]. Several of these infections resulted of contamination of respiratory equipment [76–78]. Therefore, contamination with this genus should not be routinely neglected.

Several clinical and metagenomic studies have described Gram-positive bacteria as a highly frequent colonizer of the skin [52,79]. The skin-associated genera *Staphylococcus*, *Propionibacterium*, *Streptococcus*, and *Corynebacterium*, were found in high abundance in surfaces frequently touched by hands of healthcare workers (HCW) such as, in order of decreasing abundance, computers, door handles, medical records, monitors, and mobiles (Fig. 3A). Most of the bacteria of these genera are harmless but may become opportunistic pathogens for immunocompromised patients [39]. Moreover, a high abundance of these genera was also observed in ventilators and pumps, suggesting that skin contact may be an essential source of contamination. *Staphylococcus* was found in all samples with a total abundance of 6% for both ICU and NICU. In the boxes area, this genus was more present on mattresses in ICU and ventilators in NICU (Fig. 3A). *Staphylococcus* can be found on the skin or in the nose of healthy patients causing no disease or only minor skin infections. However, several species can be deadly when invading bloodstream, joints, bones, lungs or heart [80]. *Staphylococcus aureus* is the most pathogenic and well-established species in the hospital environment.

Furthermore, coagulase-negative species such as *S. epidermidis*, *S. sciuri*, and *S. haemolyticus* are also an emerging problem in UTIs [80–82]. *Streptococcus* was found in all samples with a total abundance of 1.7% and 5% for ICU and NICU, respectively. In the boxes area, this genus was more present on monitors for both ICU and NICU (Fig. 3Aa). Nosocomial infections with *Streptococcus spp.* are often associated with respiratory or skin diseases [83] and cause long days of hospitalization [84]. Species such as *S. pneumoniae* is the first most common cause of fatal bacterial pneumonia in developing countries with high morbidity in children [85]. Group B *Streptococcus*, a commensal bacterium, is the leading cause of death from early-onset infections in the neonate [86].

Other Gram-positive genera related to nosocomial infection were found but in intermediate abundance (0.5-1%), as, e.g., *Gemella*, *Enterococcus*, and *Clostridium*. These genera were present mainly in NICU being more abundant in ventilators (Fig. 3Aa). However, they were also found in pumps (*Enterococcus)*, door handles (*Clostridium*) and computers (*Gemella*). A very low abundance (≤0.1%) of these genera was observed in ICU samples, e.g., room cards).

### Gram-negative bacteria

In the last decade, Gram-negative strains have gotten attention in their ability to spread their antibiotic resistance in hospital environments [87]. New molecular protocols, such as NGS, have allowed identifying emerging threats associated with nosocomial infections and multidrug-resistant [16]. An analysis involving eight ICUs reported that Gram-negative organisms were the principal responsible for HAI [88]. Here, the five more abundant Gram-negative genera found in ICU samples were, in order of decreasing abundance, *Pseudomonas*, *Pseudoxanthomonas*, *Castellaniella*, *Acinetobacter*, *Thermomonas*. Whereas for NICU were *Castellaniella*, *Delftia*, *Acinetobacter*, *Stenotrophomonas*, *Pseudomonas*. Previous studies have reported *Acinetobacter* and *Pseudomonas* as typical Gram-negative genera on (N)ICU surfaces [39].

*Pseudomonas*, *Acinetobacter*, *Delftia*, and *Stenotrophomonas* are known for their facultative pathogenic nature and for being nosocomial bacteria. *Pseudomonas* constituted 5%, and 2.6% of the bacterial community at ICU and NICU, respectively. The ICU mattresses and NICU drug station contained the highest abundance (Fig. 3A). *Acinetobacter* was found in all samples with a total abundance of 4% and 3% for ICU and NICU, respectively. *Acinetobacter* was more frequent on ICU mobiles. In the boxes area, this genus was more present on ICU mattresses and NICU monitors (Fig. 3A). *Delftia* showed an abundance of 1% at ICU and 5.5% at NICU, being more frequent on mobiles and NICU monitors (Fig. 3A). *Stenotrophomonas* showed an abundance of 1.2% and 2.7% for ICU and NICU, respectively. This genus was more present on NICU drug station and ventilators (Fig. 3A). *Enterobacter* and *Escherichia*_*Shigella* were also found in high abundance in both units, mainly in pumps (NICU), and ICU bed matrasses and rails (Fig. 3Aa). In general, fecal indicators (*Enterobacter, Escherichia, Bacteroides, Anaerobiospirillum, and Parabacteroides*) were more frequent in bed matrasses and rails for ICU, and pumps for NICU (Fig. S3Aa).

Others Gram-negative HAI-related genera such as *Elizabethkingia* (bed rails), *Neisseria* (mobiles), *Haemophilus* (ventilators), *Leptotrichia* (mobiles), and *Serratia* (pumps) were primarily present only in NICU. *Prevotella* and *Sphingomonas* were found in both units in intermediate abundance.

